# Modelling variability in functional brain networks using embeddings

**DOI:** 10.1101/2024.01.29.577718

**Authors:** Rukuang Huang, Chetan Gohil, Mark Woolrich

## Abstract

Functional neuroimaging techniques allow us to estimate functional networks that underlie cognition. However, these functional networks are often estimated at the group level and do not allow for the discovery of, nor benefit from, subpopulation structure in the data, i.e. the fact that some recording sessions maybe more similar than others. Here, we propose the use of embedding vectors (c.f. word embedding in Natural Language Processing) to explicitly model individual sessions while inferring dynamic networks across a group. This vector is effectively a “fingerprint” for each session, which can cluster sessions with similar functional networks together in a learnt embedding space. We apply this approach to estimate dynamic functional connectivity, using Hidden Markov Models (HMMs), which are popular methods for inferring dynamic networks, to model individual sessions in neuroimaging data. We call this approach HIVE (HMM with Integrated Variability Estimation). Using simulated data, we show that HIVE can recover the true, underlying inter-session variability and show improved performance over existing approaches. Using real magnetoencephalography data, we show the learnt embedding vectors (session fingerprints) reflect meaningful sources of variation across a population (demographics, scanner types, sites, etc). Overall, HIVE provides a powerful new technique for modelling individual sessions while leveraging information available across an entire group.

**Highlights:**

- We proposed the use of embedding vectors and a novel variability encoding block for inferring individualised brain networks in neuroimaging data.
- We apply this approach to estimate dynamic functional connectivity using the Hidden Markov Models (HMMs) and explicitly model variability in the training dataset. We call this new model HIVE (HMM with Integrated Variability Estimation)
- We demonstrate the advantages of HIVE over traditional approaches using both simulated and real MEG data.
- We show HIVE learns meaningful variability in the data (e.g. measurement site, scanner type, demographics) in an unsupervised manner.
- The datasets and scripts for performing all the analysis in this paper are made publicly available.

## 1 Introduction

A number of studies have shown that the brain recruits functional networks during task ([21, 40, 36, 14]), and at rest ([4, 9, 17]). A measure of functional networks is *functional connectivity* (**FC**), defined as the temporal correlation between spatially remote regions ([22]).

Much of these studies has used time-averaged, or static, estimates ([9, 30, 20, 19, 6]). However, given the dynamic nature of brain activity, it is increasingly important to understand how FC between distinct brain regions fluctuates over time. Dynamic FC methods include the sliding window approach combined with clustering ([1, 11]), the Hidden Markov Models (HMMs, [3, 48]), and the recently developed Dynamic Network Modes (DyNeMo, [25]).

A common assumption in both static and dynamic FC methods, is that the same network, or set of networks, are shared by all recording sessions. This is an unrealistic assumption as functional brain activity is known to possess significant heterogeneity related to subject demographics, scanner types, sites, etc. As a result, the accurate estimation of subject/session-specific networks is paramount for characterising differences between individuals and for downstream prediction tasks.

Session-specific estimates of static FC can be easily obtained, or by using post-hoc analyses of model-based dynamic FC. For example, session-specific estimates can be obtained by combining HMMs with *dual estimation*, which has a similar rationale as dual regression in Independent Component Analysis (ICA, [38], [5]). However, these approaches do not typically allow the inference of dynamic networks from the data to benefit from the subpopulation structure within the group. An alternative to overcome this is to use hierarchical models that capture variability across sessions. For example, PROFUMO ([28, 27]) has been successfully used to capture aspects of subject variability in functional modes in fMRI data. However, variability within a population is typically modelled using very simple parametric distributions (e.g. univariate Gaussian), which fails to capture the rich, multivariate variability expected across sessions.

In this paper, we employ embedding vectors, a technique widely used in the Natural Language Processing literature for characterising semantic relationships between words ([37]), to explicitly model variability in the basis set of networks used to describe the training data sessions. Embedding vectors have been previously used to deal with between-subject variability in the context of supervised learning ([15, 12]). Here, we show how we can incorporate embedding vectors into a generative model of FC to perform unsupervised learning, and to model variability of covariance matrices that describe network activity. We apply this approach to the generative model of the HMMs ([3, 48, 46, 49, 29, 40]), and call this new model **HIVE** (HMM with Integrated Variability Estimation).

Using simulated and real MEG datasets, we show that HIVE can infer the underlying population structure in an unsupervised manner. This helps to give a more accurate description of the session-specific network estimates than traditional method based on the group-level model. While we focus here on the estimation of dynamic FC using the HMM on MEG data, this approach can be readily adapted to other methods that estimate static or dynamic FC, and on other modalities including EEG and fMRI. Source code for the HIVE model is available in the *osl-dynamics* toolbox ([24]) and scripts to reproduce results in this paper are available here: github.com/OHBA-analysis/Huang2024_ModelVariabilityWithEmbeddings.

## 2 Methods

We start by describing how embedding vectors can be incorporated into a generative model of FC to describe variability in network activity.

### 2.1 The use of embedding vectors

The use of word embeddings is a common practice in Natural language processing (NLP, [37]). A word embedding is a real-valued vector that encodes the meaning of the word such that words closer in the vector space have similar meanings. In Figure 1a, we show an example of word embeddings. Here the x-axis encodes whether a word is for animals or non-animals and the y-axis encodes whether the object can fly or not. Similarly, we can assign a vector for each recording session and hope that sessions with similar properties are close to each other in the vector space. In Figure 1b, we show an illustration of possible variability captured by embedding vectors. Here the x-axis encodes the direction of increasing beta power in motor network whereas the y-axis encodes the direction of increasing peak alpha frequency in visual network.

**Figure 1:**
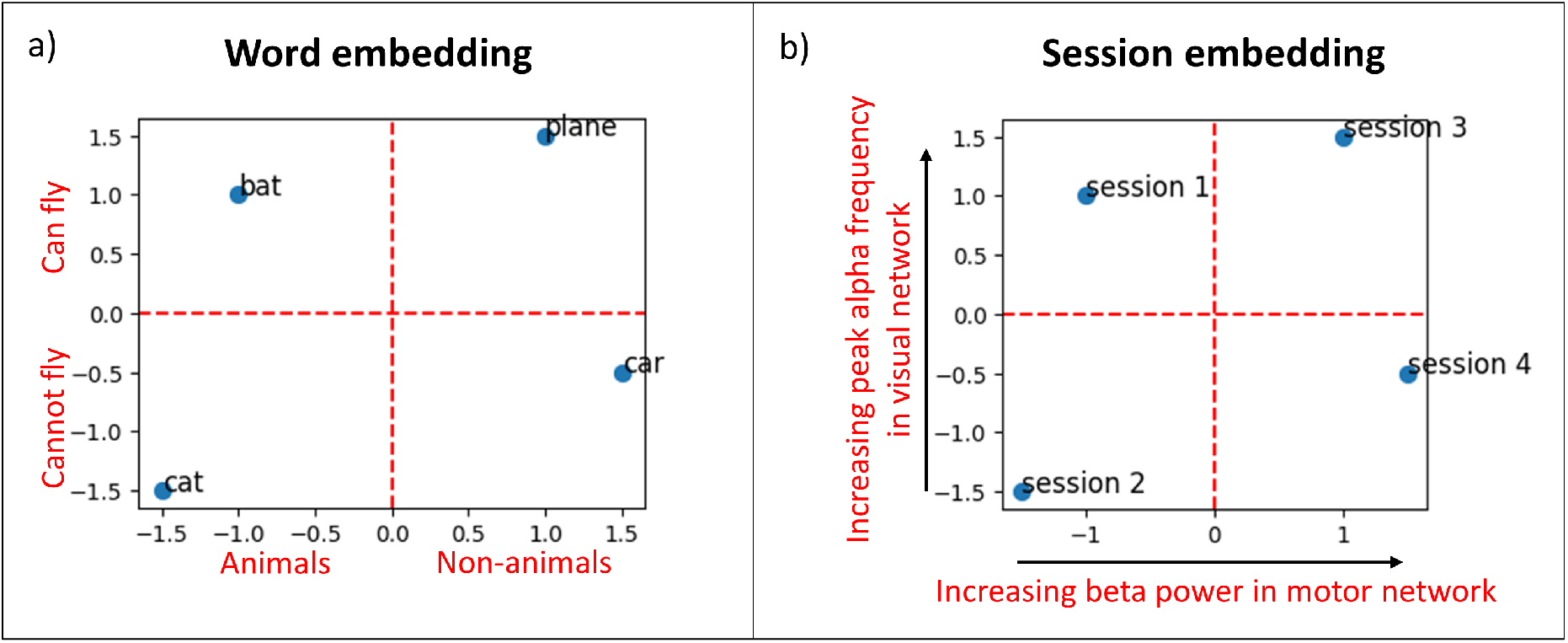
Examples of embedding vectors. a) Example of word embedding vectors. The x-axis encodes whether a word is for animals or non-animals and the y-axis encodes whether the object can fly or not. b) Illustration of possible variability captured by embedding vectors in brain networks found in electrophysiological data such as M/EEG. The x-axis encodes the direction of increasing beta power in motor network whereas the y-axis encodes the direction of increasing peak alpha frequency in visual network.

### 2.2 Generating session-specific covariances

In this section we outline the process of modelling session-specific covariances. For *n* ∈ ℕ, let [*n*] be the set {1, …, *n*}. Suppose {***D***_*j*_ : *j* ∈ [*J*]} is the set of covariances that describes group-level functional networks, where *J* is called the *number of states*. Subsequently, we assume there are *N* sessions in the dataset and each session *i* ∈ [*N*] has its own set of covariances 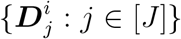. We want the covariances for different sessions to respect (be regularised) by the relationships between sessions in the population. Furthermore, we want the session-specific covariances to vary around the group-level covariances. This is done using our novel variability encoding block illustrated in Figure 2.

**Figure 2:**
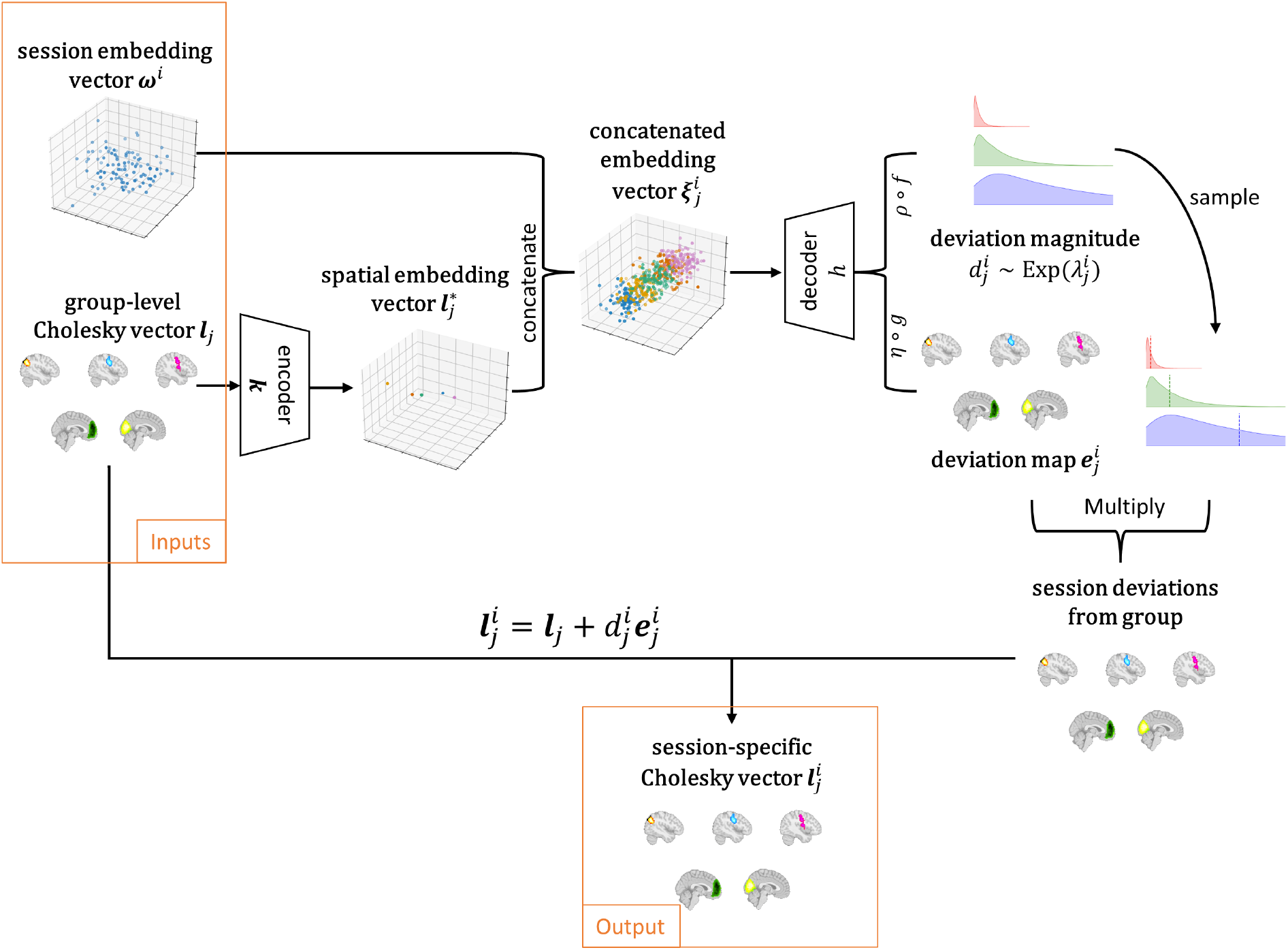
Variability encoding block. Session-specific information from the session embedding vectors ***ω***^*i*^ and state-specific information from the group-level Cholesky vectors ***l***_*j*_ are combined to generate session and state-specific covariances 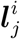. Here the **encoder** *k* is an affine transformation that condenses state-specific information to a lower dimensional space. The **decoder** *h* is a multi-layer perceptron (MLP) that decodes state and session-specific deviations from the concatenated embedding vectors 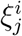.

Given a session *i* and a state *j*, we want to generate the state and session-specific covariance matrix 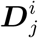. Let ***ω***^*i*^ be the session embedding vector of session *i*, which is a vector of pre-defined length *n*_*ω*_ that provides a “fingerprint” for this session. Next, to ensure legitimacy of generated session-specific covariances, we work in the Cholesky space. Let 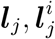 be the flattened vectors of the lower triangular entries of the Cholesky factors 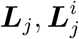 of 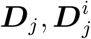 respectively, then the session-specific Cholesky vector 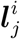 is generated by adding a deviation to the group-level Cholesky vector ***l***_*j*_ with

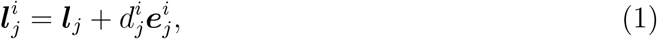

where 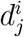 is a scalar representing the spatially global magnitude of the deviation and 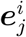 is a vector representing the standardised pattern of the spatial deviation in the sense that it has unit sample standard deviation. We would expect the deviation magnitude 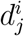 to be small so that the session-specific covariances vary around the group-level covariances, while enforcing positivity of this quantity. Hence we put an exponential prior on 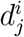 :

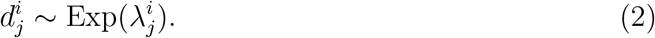

The rate parameter 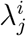 and deviation pattern 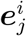 are generated through

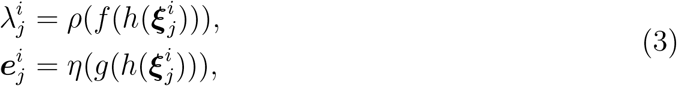

where *ρ* is the softplus function to ensure positivity of 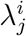 and *η* is a Layer Normalisation layer ([2]) with non-trainable scale parameter of 1 to ensure unit standard deviation of ***e***^*i*^. Affine functions *f, g* are learnable transformations that extract different information from the hidden state given by the decoder *h*, a multi-layer perceptron (MLP, [39]). 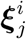 is the key object called a **concatenated embedding vector** that encodes both session and state-specific information. It is formed by concatenating the session embedding vector ***ω***^*i*^ and spatial embedding vector 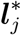, which is a lower dimensional representation of the Cholesky vectors:

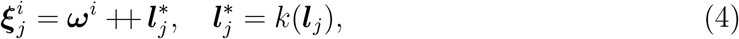

where *k* is an affine transformation that serves as an encoder to encode spatial information and ++ is a binary operator that concatenates two vectors.

### 2.3 Generative model of HIVE

We next consider how we can use our proposed variability encoding block as part of a method for estimating functional networks, by using the example of the HMM. The HMM estimates networks and their dynamics by partitioning time series data into a finite number of states, where each state has a specific spatial-temporal pattern ([3, 48]). The generative model of HMM describes how the time series data is generated given the HMM model parameters.

Let ***x***_*t*_ be the data we observe at time *t* ∈ [*T*]. Here *T* is the total number of samples/time points and ***x***_*t*_ is a vector of length *N*_*c*_ - the number of channels. In HMM, it is assumed that the data are generated by

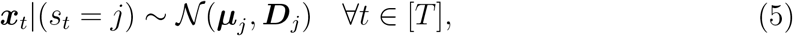

where ***µ***_*j*_, ***D***_*j*_ are state-specific means and covariances that describe the spatiospectral pattern for each state *j* ∈ [*J*]. In HMM, the process {*s*_*t*_ : *t* ∈ [*T*]*}* follows a discrete homogeneous Markov process ([23]). i.e.

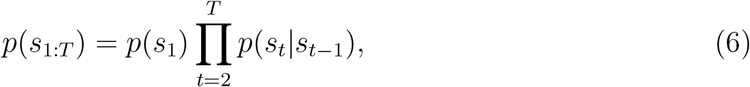

where *p*(*s*_1:*T*_) is a shorthand for the joint distribution *p*(*s*_1_, …, *s*_*T*_), *p*(*s*_*t*_ = *j*|*s*_*t*−1_ = *j*^*′*^) = *A*_*j*_*′*_*j*_ does not depend on time and is the (*j′, j*)-th entry to the transition probability matrix ***A***.

For simplicity, in HIVE, we assume that the state-specific means are fixed to zero (i.e. ***µ***_*j*_ = 0 ∀*j* ∈ [*J*]). Suppose for each session *i*, the data 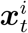 is generated by

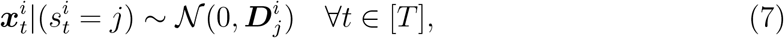

where 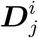 is generated through the variability encoding block described in Section 2.2. It should be noted that the variability encoding block can be easily adapted to generate session-specific means 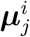 by excluding the transformation into Cholesky space. We further assume all sessions have the same transition probability matrix, i.e. ***A***^*i*^ = ***A*** ∀*i* ∈ [*N*]. The complete generative process of HIVE is summarised in Figure 3.

**Figure 3:**
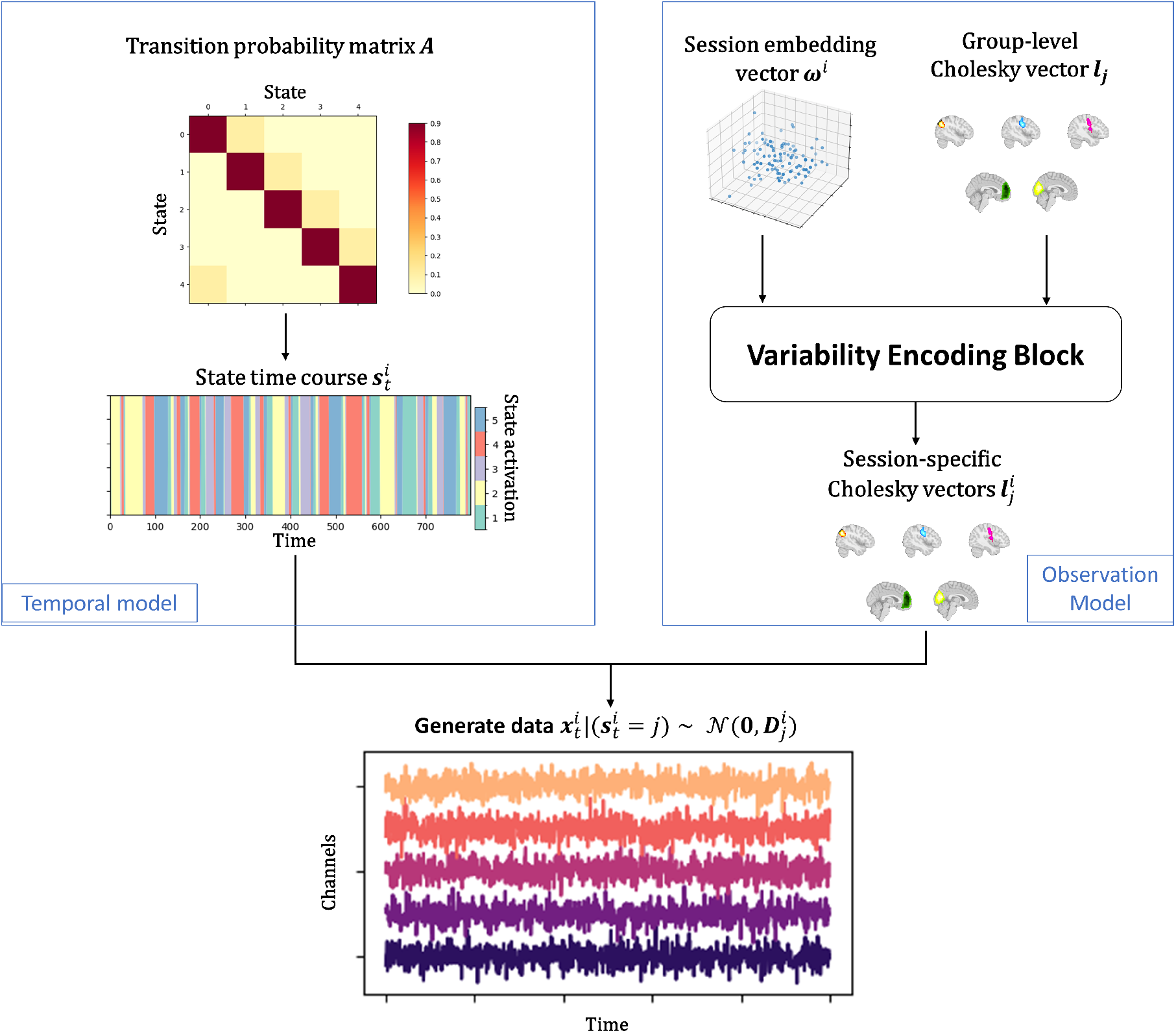
Generative model of HIVE. State time courses are generated from the group-level transition probability matrix ***A*** through a Markov process. At the same time, session embedding vectors ***ω***^*i*^ and group-level Cholesky vectors ***l***_*j*_ are passed to the variability encoding block to give session-specific Cholesky vectors 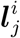. Given the state 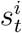 at each time, the observed data is generated through the multivariate Gaussian likelihood.

### 2.4 Dual estimation in the traditional HMM approach

The state means and covariances ***µ***_*j*_, ***D***_*j*_ are group-level estimates. Similar to dual regression in ICA, dual estimation ([47]) is a way to compute session-specific means and covariances in HMM. For each session, we fix the inferred state probabilities from the group-level HMM and re-estimate the observation model parameters with the session’s data (See A.1 for details). In this paper, we use **HMM-DE** (HMM with dual estimation) to refer to the pipeline of training a group-level HMM followed by dual estimation for estimating session-specific parameters (which correspond to the session-specific networks).

### 2.5 Inference

The parameters of HIVE include the group-level transition probability matrix ***A***, the embedding vectors ***ω*** = {***ω***^*i*^ : *i* ∈ [*N*]}, the group-level Cholesky vector ***l*** = {***l***_*j*_ : *j* ∈ [*J*]}, as well as the weights of the learnable transformations *h, f, g, k*. To perform inference on these parameters, we employ the variational expectation–maximization (EM) algorithm ([18]), in particular, a variant of the Baum-Welch algorithm ([52]). During the “E-step”, we update the state probabilities (the posterior distribution of the states). During the “M-step”, given the state probabilities, we update the transition probability matrix ***A*** with the stochastic update technique used in [45] and all other parameters by minimising the objective function (variational free energy given the state probabilities, [7])

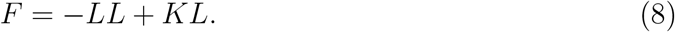

The first term is the negative log-likelihood term. Let *T*_*i*_ be the number of samples for session 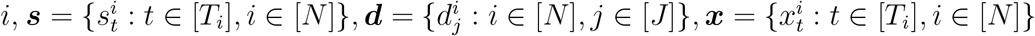, and *q*(***s***), *q*(***d***) be the posterior distribution on ***s*** and ***d*** respectively. Denote 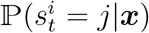 the posterior probability that for session *i*, the state at time *t* is *j*, we have

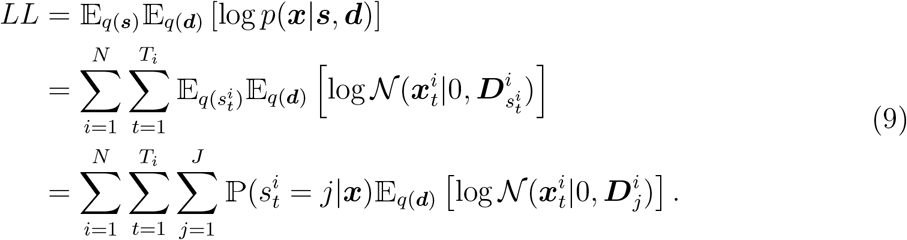

Notice we omit the dependence of 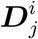 on ***d, ω, l*** and weights of *h, f, g, k* for readability. The second term is the KL loss term on the observation model due to the deviation magnitudes,

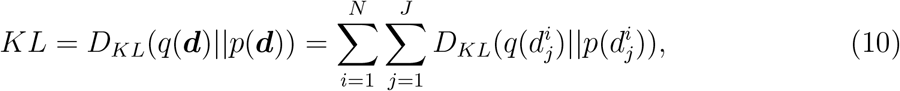

where *D*_*KL*_(*q*(.)||*p*(.)) is the Kullback–Leibler divergence ([35]) and we approximate the posterior distribution *p*(***d***|***x***) with a mean field approximation:

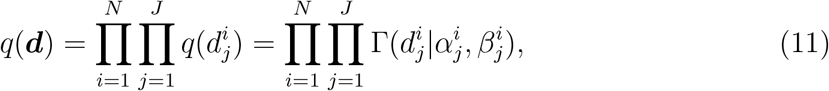

where the shape 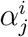 and the rate 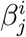 parameters of the Gamma distributions Γ are learnable parameters. We can write down the analytic solution for *KL* and its gradients, and we used the reparameterisation trick ([34]) to get an estimate of the gradient of *LL* with respect to 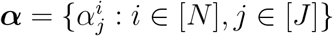 and 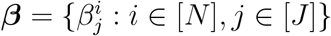. During training, we minimise *F* with the Adam optimiser ([33]). The complete inference framework including forward pass and backward update is summarised in Figure 4.

**Figure 4:**
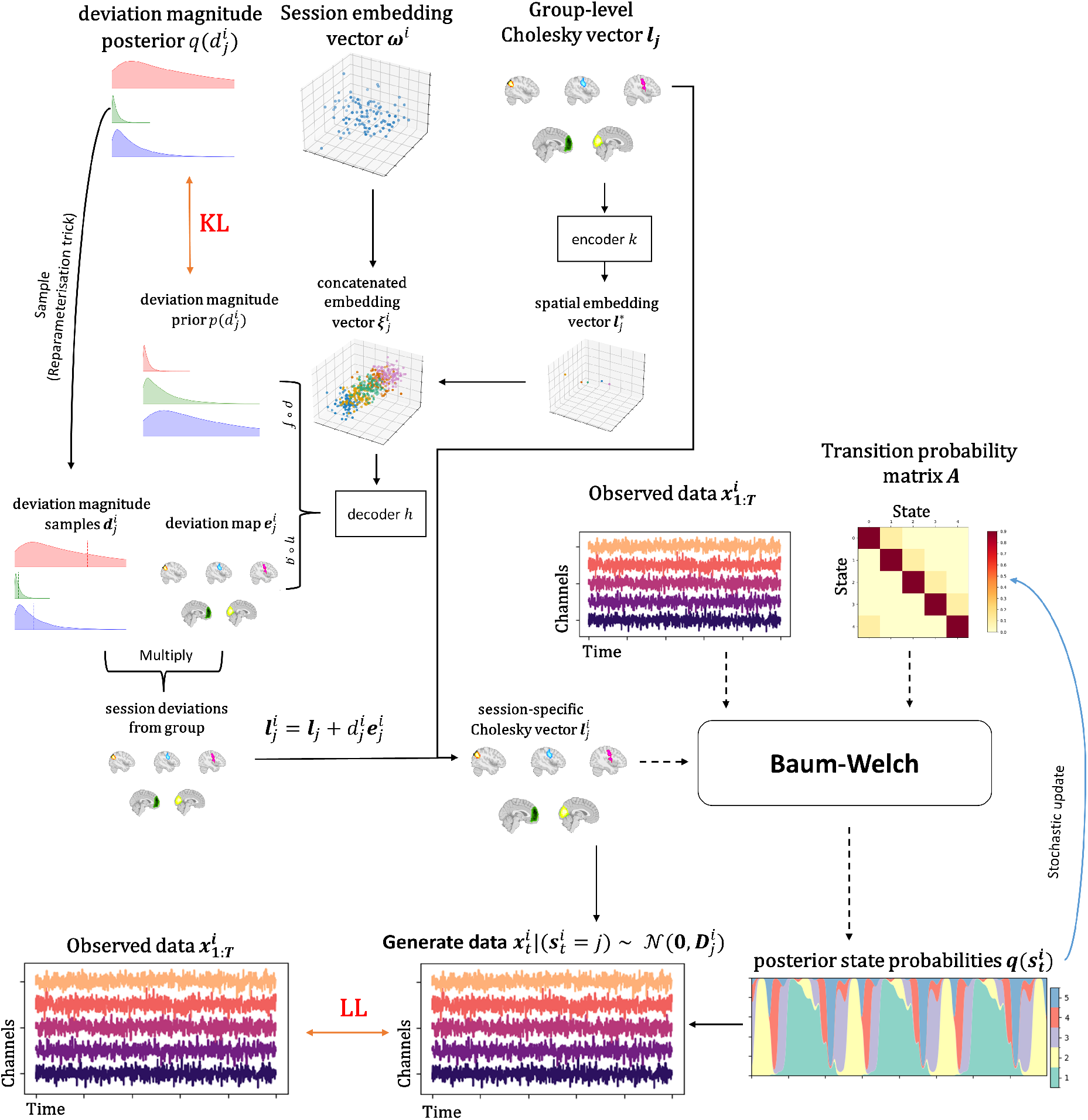
The full HIVE inference framework. The session embedding vector ***ω***^*i*^, group-level Cholesky vector ***l***_*j*_ are used to generate the deviation map 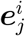 and the deviation magnitude prior distribution 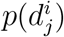. We sample 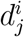 from the deviation magnitude posterior distribution 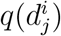 using the reparameterisation trick and use the drawn samples, which give the session-specific Cholesky vector 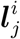, together with the deviation map 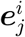 and group-level Cholesky vector ***l***_*j*_. The prior and posterior distributions 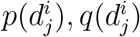 are used to calculate the *KL* term in the loss function. Subsequently, we get the posterior state probabilities 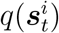 with the Baum-Welch algorithm which takes the observed data 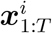, the group-level transition probability matrix ***A*** and the session-specific Cholesky vectors 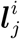 as inputs. Finally, we calculate the *LL* term in the loss function with the observed data 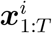. The black solid arrows in the plot show connections where gradient flows through during back-propagation and the dashed black arrows show connections where gradient does not flow through. The blue arrow shows the stochastic update step for the group-level transition probability matrix ***A***.

#### Choosing the embedding dimension *n*_*ω*_

We need to pre-define the length *n*_*ω*_ of the embedding vectors ***ω***^*i*^. A smaller *n*_*ω*_ will lead to a more parsimonious model but there could be a risk of over-regularisation. On the other hand, a larger *n*_*ω*_ will lead to a more flexible model but could lead to overfitting. In practice, we use the following strategy. Firstly, we define a set of candidate values for *n*_*ω*_, say 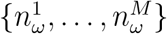 where 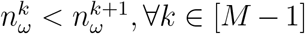. Then we train the model with each of these values a number of times independently and for each run, we get the variational free energy of the model on the training data. Starting from *k* = 1, we would then do progressive t-tests with the alternative hypothesis that the variational free energy of the model with 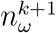 is lower than the variational free energy of the model with 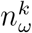. We would stop when the null hypothesis is not rejected and choose the model with 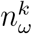 as the final model, or the null hypothesis is never rejected until the largest candidate 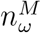 is reached, in which case we choose the model with 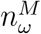 as the final model. See Section A.6 for details.

#### Initialisation and annealing during training

A good initialisation is paramount in training deep neural networks to avoid slow and unstable training. The variational parameters ***α, β*** are initialised by calculating the session-specific variations of static covariance of the data (see A.2 for details).

During training, we employed two annealing techniques. The first one is KL annealing ([8]) where we start the training without the KL term in the loss and gradually increase the contribution of the KL term, which is a common practice for training recurrent neural network-based variational autoencoders ([8]). The second one is that we anneal the sampling process of the variational distribution of deviation magnitudes *q*(***d***) when applying the reparemeterisation trick:

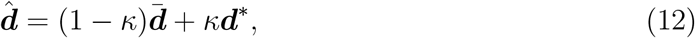

where 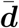 is the expectation of *q*(***d***), ***d***^∗^ is a sample from *q*(***d***) and *κ* is the annealing factor that is zero at the beginning of the training and gradually increases to one. We find this significantly improves the convergence of the model and we believe this serves as an exploration-exploitation mechanism, which is extensively studied in the fields of reinforcement learning ([51]) and Bayesian Optimisation ([31]). At the beginning of training, only the expectation is used and the gradient will intend to push the mean deviations to the correct position. This is the exploration phase. As training progresses, we gradually take into account the variance and higher moments of the variational distribution, which allows the training to fine tune the variational parameters. This is the exploitation phase.

### 2.6 Datasets

#### 2.6.1 Simulated datasets

In the simulation studies, we simulate data with an HMM. To do this, we specify a transition probability matrix and randomly simulate covariance matrices for each state (sessions can have different covariances). Subsequently, a Markov chain is simulated with the pre-specified transition probability matrix. Lastly, at each time point, data is simulated using a multivariate Gaussian distribution with the state covariance matrix which is active at the given time. Gaussian noise is also added to the final time series data. The simulated data for the 3 simulation studies are described below.

- **Simulation 1**. We simulate data with *N* = 10, *T*_*i*_ = 25, 600, ∀*i* ∈ [*N*], *N*_*c*_ = 11, *J* = 5. Here the state covariance matrices of sessions 1 and 2 are altered and all other sessions have the same unaltered group-level covariances - session 1 has increased variance in channel 1 and decrease variance in channel 2 whilst session 2 has decreased variance in channel 1 and increased variance in channel 2. For both session 1 and 2, the increased variances are 5 times and the decreased variances are 20% of the group-level variance.
- **Simulation 2**. Here *N* = 100, *T*_*i*_ = 3, 000, ∀*i* ∈ [*N*], *N*_*c*_ = 40, *J* = 5. Sessions are assigned into 3 groups and an embedding vector for each session is simulated according to the session’s assigned group. Session-specific covariances are simulated based on the simulated embedding vectors. (See A.3 for details).
- **Simulation 3**. In this study 4 datasets are simulated with *N* ∈ {5, 10, 50, 100*}* and *T*_*i*_ = 3000, ∀*i* ∈ [*N*], *N*_*c*_ = 40, *J* = 5. Session-specific covariances are simulated in the same way as in Simulation 2.

#### 2.6.2 Real MEG data

We demonstrate the use cases of the proposed model with 3 publicly available MEG datasets, including two resting-state and one visual task dataset. The datasets are source-reconstructed and parcellated to 38 regions of interest. We describe the steps of data processing before model training below.

- **Raw data**. The first resting-state dataset ([43], we refer to this dataset as the **Cam-CAN** dataset) contains eyes-closed data from 612 healthy participants. These data were collected using an Elekta Neuromag Vectorview 306 scanner at a sampling frequency of 1kHz. A highpass filter of 0.03Hz and MaxFilter were applied. In the visual task MEG dataset ([50], we refer to this dataset as the **Wakeman-Henson** dataset), each of the 19 health participants were scanned 6 times, during which 3 types of visual stimuli were shown to the participants. The data were also collected using an Elekta Neuromag Vectorview 306 scanner. The second resting-state dataset was collected suing a 275-channel CTF scanner. This dataset (we refer to this dataset as the **Nottingham** dataset) contains eyes-closed data from 64 healthy participants, collected at Nottingham University, UK as part of the MEGUK partnership.
- **Preprocessing**. The Cam-CAN and Nottingham datasets were preprocessed with the same pipeline using the OHBA software library (OSL). The data were band-pass filtered between 0.5Hz and 125Hz, followed by a notch filter at 50Hz and 100Hz. Then the data were downsampled to 250Hz before automated bad segment and bad channel detection were applied to remove abnormally noisy segments and channels of the recording. Finally, an independent component analysis (ICA) with 64 components was applied to identify artifacts. The preprocessing of the Wakeman-Henson dataset is the same as above except that in the end, ICA with 40 components was applied.
- **Source reconstruction and parcellation**. Coregistration and source reconstruction were done using OSL. Structural data were coregistered with the MEG data using an iterative close-point algorithm and digitised head points acquired with a Polhemous pen were matched to individual subject’s scalp surfaces extracted with FSL’s BET tool ([32], [42]). The nose was not included in the coregistration as the structural MRI images were defaced. Preprocessed sensor data were source reconstructed onto an 8mm isotropic grid using a linearly constrained minimum variance beamformer ([44]). Voxels were then parcellated into 38 anatomically defined regions of interest, before the symmetric spatial leakage correction described in [13] was applied.
- **Data preparation**. Before model training, we follow the preparation steps described in [25]. The data were time-delay embedded with *±*7 lags. Then principal component analysis (PCA) was used to reduce the dimensionality to 80 channels before a standardisation step (z-transform) was applied to make sure each channel has zero mean and variance of one.

## 3 Results

### 3.1 Simulation

Here we show the results of training HIVE on the 3 simulated datasets described in Section 2.6.1 and compare these results with those from HMM-DE.

#### Simulation 1: Variability encoding block learns multivariate session-specific covariance deviations

We aim to test if patterns of deviations across multiple channels can be learnt by the generative model, in particular, the variability encoding block, during inference. This is an important feature in the sense that we want the generative model to act as a prior that regularises how each session can deviate from the group. Firstly, we can see from Figure 5a that both HMM-DE and HIVE can infer the group-level covariances correctly. What HIVE offers in addition is shown in Figure 5b where the inferred session embedding vectors are plotted. We can clearly see that the embedding vectors for sessions 1 and 2 are far away from the cluster formed by other sessions which have unaltered covariances, which matches well with the ground truth. The model’s capability to encode deviation pattern across multiple channels in the generative model is illustrated in Figure 5c, in which the simulated patterns of deviation for sessions 1 and 2 can be generated from the trained variability encoding block. Although HMM-DE can infer session-specific covariances via dual estimation, it cannot generate data with patterns that deviate from the group mean. This is because intrinsically HMM-DE is a group-level model.

**Figure 5:**
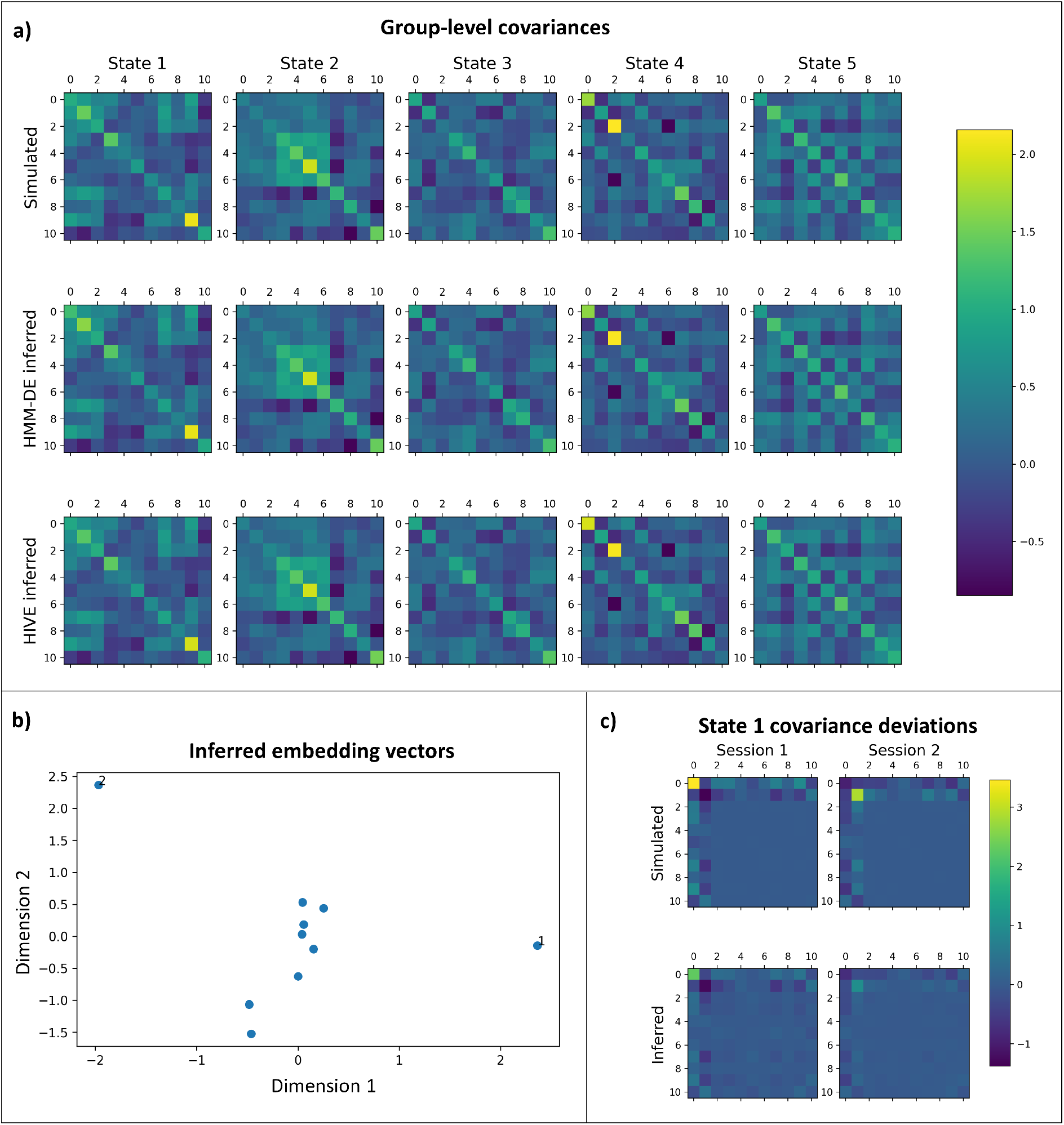
Variability encoding block learns session-specific covariance deviations. *Results obtained on Simulation 1*. a) Simulated (top row), HMM-DE inferred (middle row) and HIVE inferred (bottom row) group-level covariances. The columns show covariances of different states. b) Inferred session embedding vectors. Sessions 1 and 2 are annotated. c) Simulated deviations (top row) and deviations from trained generative model of HIVE (bottom row) for state 1 for session 1 (left column) and for session 2 (right column).

#### Simulation 2: Underlying subpopulation structure is inferred by HIVE

In this simulation study, we aim to show that HIVE can recover the ground truth underlying subpopulation structure (simulated embedding vectors of sessions). This is demonstrated in Figure 6a, where the ground truth grouping of sessions is recovered by the inferred session embedding vectors. In particular, sessions which are close together in the simulated space (e.g. sessions 7 and 45) stay close in the inferred space. Figure 6b shows both HMM-DE and HIVE can infer the state time courses perfectly. Furthermore, we can see from Figure 6c that both approaches can recover the pairwise session relationship between session-specific covariances, though HMM-DE overestimates the pairwise distance due to noise added to the data, whereas HIVE has the preferable behaviour of underestimating the pairwise distance due to the regularising effect of the prior on the deviation magnitude. This prior makes the inferred session-specific covariances more similar to the group average, when there is insufficient evidence available in the data to do otherwise.

**Figure 6:**
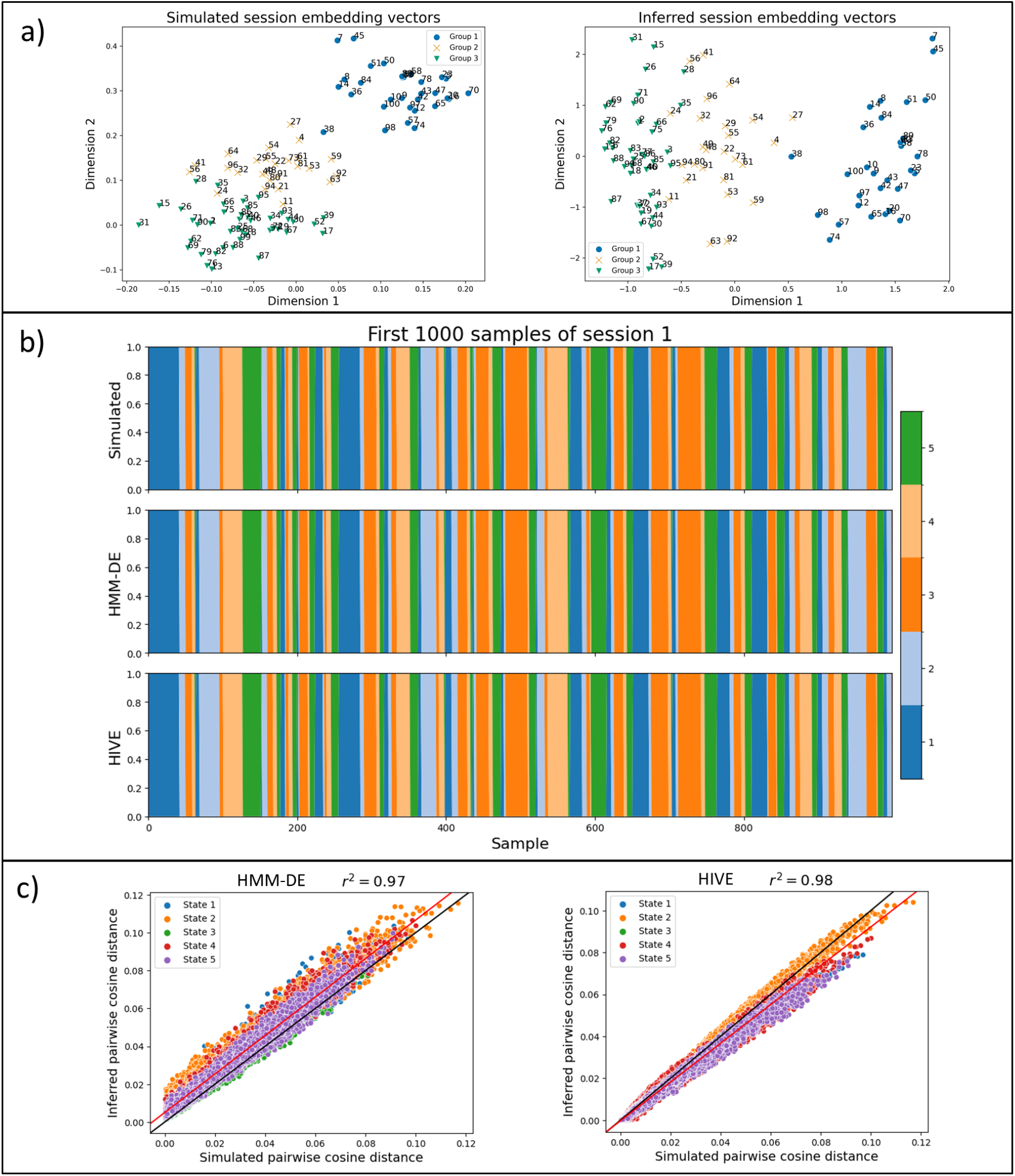
HIVE infers the underlying subpopulation structure. *Results obtained on Simulation 2*. a) Simulated (left) and inferred (right) session embedding vectors. Each point is marked and coloured by the ground truth group assignment, and is annotated by the session number. b) Simulated (top), HMM-DE inferred (middle) and HIVE inferred (bottom) state time courses. c) Session-pairwise cosine distance of simulated (x-axis) against inferred (y-axis) covariances from HMM-DE (left) and HIVE (right). The black line shows the *y* = *x* line, which corresponds to optimal performance, and the red line is a fitted line through the points, with the coefficient of determinant *r*^2^ displayed in the title.

#### Simulation 3: HIVE performs more accurate inference than HMM-DE

Now we focus on comparing HIVE and HMM-DE in terms of accuracy of inferred session-specific covariances. Both HIVE and HMM-DE are trained 10 times to get a more accurate and stable estimation of the accuracy. From Figure 7 we see HIVE can infer more accurate session-specific covariances than HMM-DE. Crucially, with the number of samples per session being kept the same, the accuracy for HIVE improves with more sessions in the dataset while that for HMM-DE does not have a significant improvement. This shows the advantage of the variability encoding block in making use of data of heterogeneous sessions to help infer on every single session.

**Figure 7:**
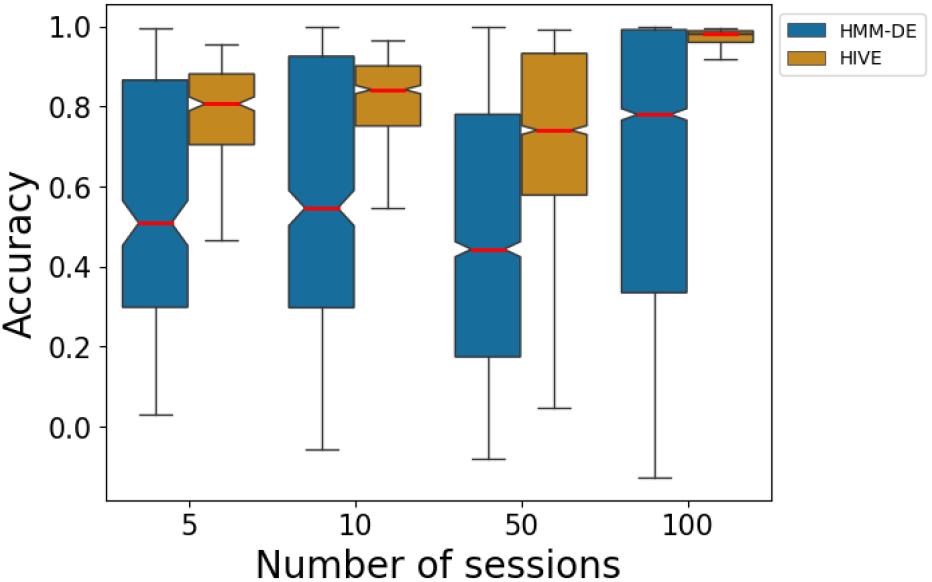
HIVE performs more accurate inference than HMM-DE. *Results obtained on Simulation 3*. Accuracy of session-specific covariances (y-axis) plotted against number of sessions in the dataset (x-axis) for HMM-DE (blue) and HIVE (orange). Here the number of samples per session is always the same. Correlation between inferred and simulated session-specific covariances is used as a metric for accuracy.

### 3.2 Real MEG data

#### 3.2.1 HIVE reveals similarities and differences between MEG recordings

In this section, we study the Wakeman-Henson dataset (see Section 2.6). This dataset contains 6 sessions for each subject. Ideally, we would expect the sessions for a subject to be more similar than sessions for different subjects. In this study, we assign each session an embedding vector and train HIVE on this data with *n*_*ω*_ = 10 (See Section A.6). The session-pairwise cosine distances of embedding vectors are plotted in Figure 8a, which shows a clear block diagonal structure - session embedding vectors from the same subject are closer together than those from different subjects. This shows the model is able to identify certain recordings which have similar deviations from the group, despite HIVE being trained in an unsupervised manner with no knowledge of which sessions belong to which subjects.

**Figure 8:**
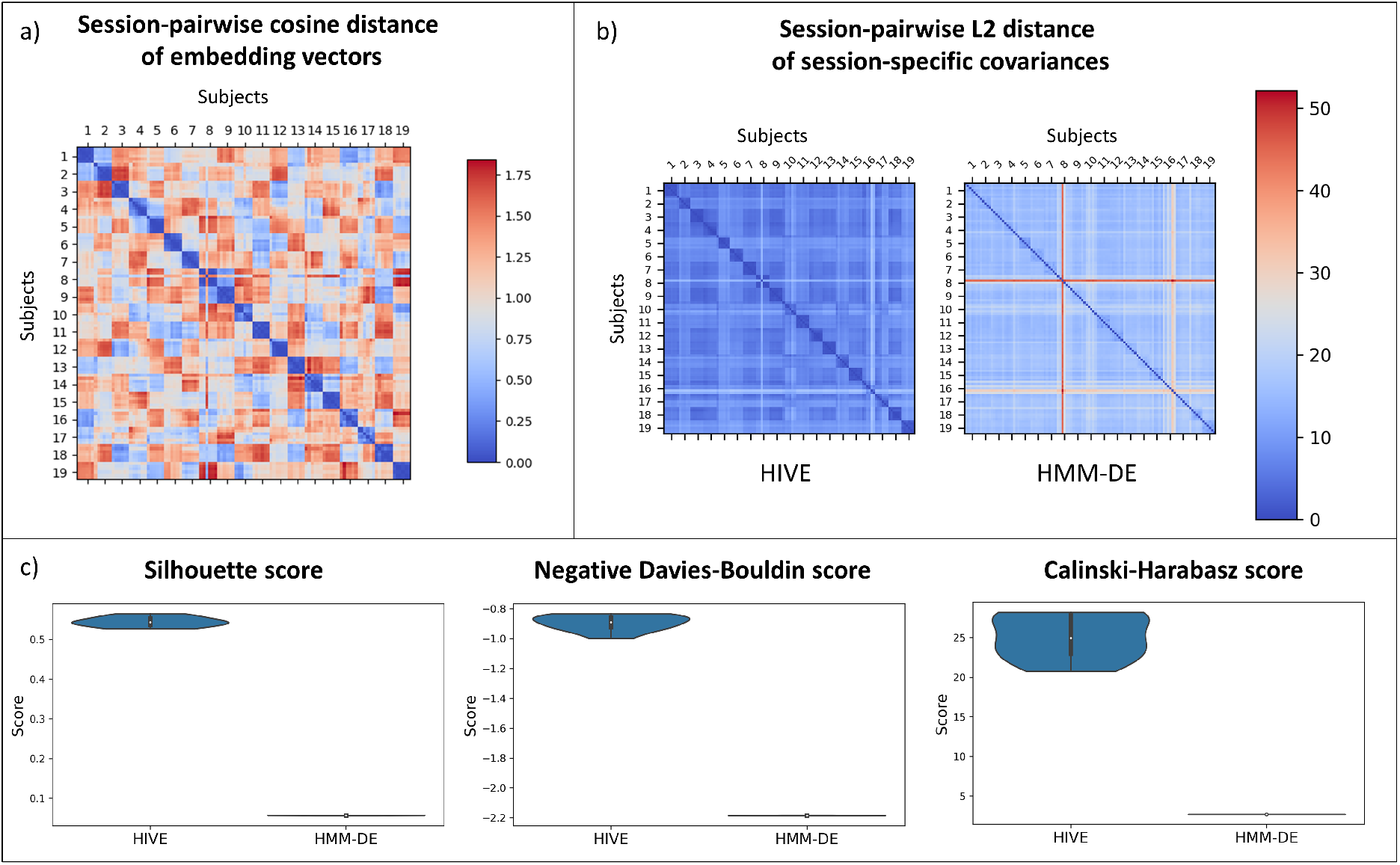
HIVE reveals similarities and differences between MEG recordings. *Results obtained on the Wakeman-Henson dataset*. a) Session-pairwise cosine distance of inferred embedding vectors. Embedding vectors for the same subject are grouped together and have smaller distance between them. b) Session-pairwise L2 distance of inferred covariances from HIVE (left) and HMM-DE (right). c) Clustering metrics - Silhouette score (left), negative Davies-Bouldin score (middle), Calinski-Harabasz score (right), based on subject labels for 10 independent runs of both approaches. Higher values for these metrics indicate better clustering.

To compare with the traditional approaches, we train both HIVE and HMM-DE on this dataset. In Figure 8b, the session-pairwise L2 distances of inferred covariances from both approaches are plotted. Although the same block diagonal structure can be seen in both approaches, it is clearer in the case of HIVE. One can observe there is a particular session (session 3 of subject 8) that has much higher distance with all other sessions, especially with HMM-DE. This is related to the fact that this particular session has very different oscillatory activity compared to other sessions (See Section A.4). In order to quantify the advantages with HIVE, we employ 3 different metrics - Silhouette score ([41]), Davies-Bouldin score ([16]) and Calinski-Harabasz score ([10]) for assessing how well the inferred covariances form distinct and well-separated subject clusters. Figure 8c shows that with all three metrics, HIVE inferred covariances form tighter and more distinct clusters than dual estimated covariances from HMM-DE.

#### 3.2.2 Embedding vectors reflect meaningful variability regarding scanner

In this study, we want to test if our model is able to differentiate data acquired by different scanners. We train on two different resting-state MEG datasets (Nottingham and Cam-CAN) described in Section 2.6. There are 128 subjects in total and 64 subjects from each datasets. To avoid the possibility that the model is biased towards either dataset, we match the age and sex profiles. Here we choose *n*_*ω*_ = 20 (See Section A.6). We can clearly see two clusters of embedding vectors inferred in Figure 9a, and at the same time an age gradient in the embedding space. This means scanner type and age information are simultaneously encoded (in different directions in the embedding space) by the embedding vectors, despite the fact that HIVE is trained unsupervised with no knowledge that the sessions were from different scanner types. We can also see a block diagonal structure in Figure 9b where subjects from the same dataset have smaller pairwise cosine distances of their inferred embedding vectors. With the clustering metrics described in Section 3.2.1, we see from Figure 9c that subject-specific covariances from HIVE form better-defined clusters based on scanners/sites.

**Figure 9:**
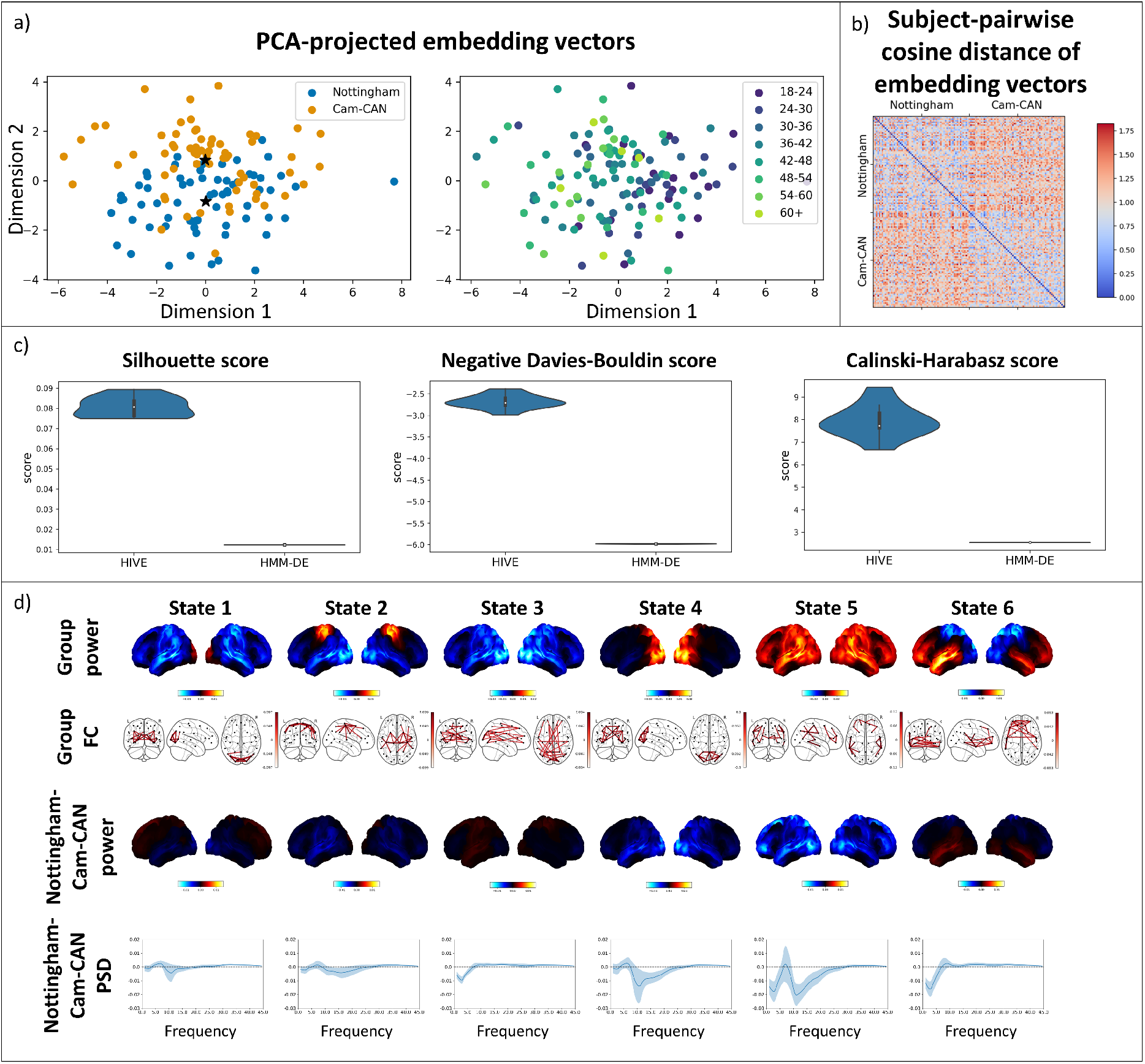
Embedding vectors reflect meaningful variability regarding scanner/site. *Results obtained with a combination of the Nottingham and Cam-CAN datasets*. a) PCA-projected embedding vectors coloured with different datasets (left, black stars indicate centroids of embedding vectors from the two datasets) and different age groups (right, older subjects are coloured with lighter colours). b) Subject-pairwise cosine distance of embedding vectors. c) Clustering metrics - Silhouette score (left), negative Davies-Bouldin score (middle), Calinski-Harabasz score (right), based on subject labels for 10 independent runs for both approaches. d) 6 states are inferred with HIVE. The first and second row shows the group level power (red areas show above average and blue areas show below average power across states) and FC maps (top 3% edges are plotted). The third row shows power difference (red areas show higher and blue areas show lower power in Nottingham subjects than Cam-CAN subjects) for each state between centroids of both datasets. The bottom row shows the difference in PSDs (solid lines show the means and shaded areas show one standard deviation across the channels) across frequencies between datasets.

HIVE also provides a way to summarise the differences in state-specific spectral content between scanner types. Ten nearest neighbours in the embedding space are selected for each of the dataset centroids and are used to give a representation of spectral contents of both datasets. In Figure 9d, we can see these differences between the “centroids” of the datasets. In particular, subjects from the Nottingham dataset generally have lower power than those from Cam-CAN, especially in posterior regions and alpha band.

#### 3.2.3 HIVE infers age structure in the population more accurately than dual estimation

Here we use the Cam-CAN dataset described in Section 2.6, that consists of 612 healthy subjects aged between 18-88 and choose *n*_*ω*_ = 50 (See Section A.6). From Figure 10a, We can clearly see an age gradient from darker to lighter dots in both plots. This means after training, age information is encoded by the embedding vectors of the subjects, despite the fact that HIVE is trained unsupervised with no knowledge of the subjects’ ages. In order to see if the learnt hidden representation given by the embedding vectors helps improve the model’s ability to distinguish between subject demographics, we try to predict age with the inferred subject-specific covariances. Here HMM-DE and HIVE with increasing embedding dimensions are trained on this dataset. Shown in Figure 10b are the distributions of accuracy (given by different folds of cross validation) of predicting age with inferred subject-specific covariances (See A.5 for details). The coefficient of determinant (*r*^2^) is used as a measure of prediction accuracy and it gradually increases with the embedding dimension until there is a significant improvement over HMM-DE when using an embedding dimension of 50. There is a slight drop in accuracy when using an embedding dimension of 100, which could be due to overfitting. Notice that when selecting the embedding dimension, we do not see a significant decrease in free energy if we increase *n*_*ω*_ from 50 to 100 (Figure 12).

**Figure 10:**
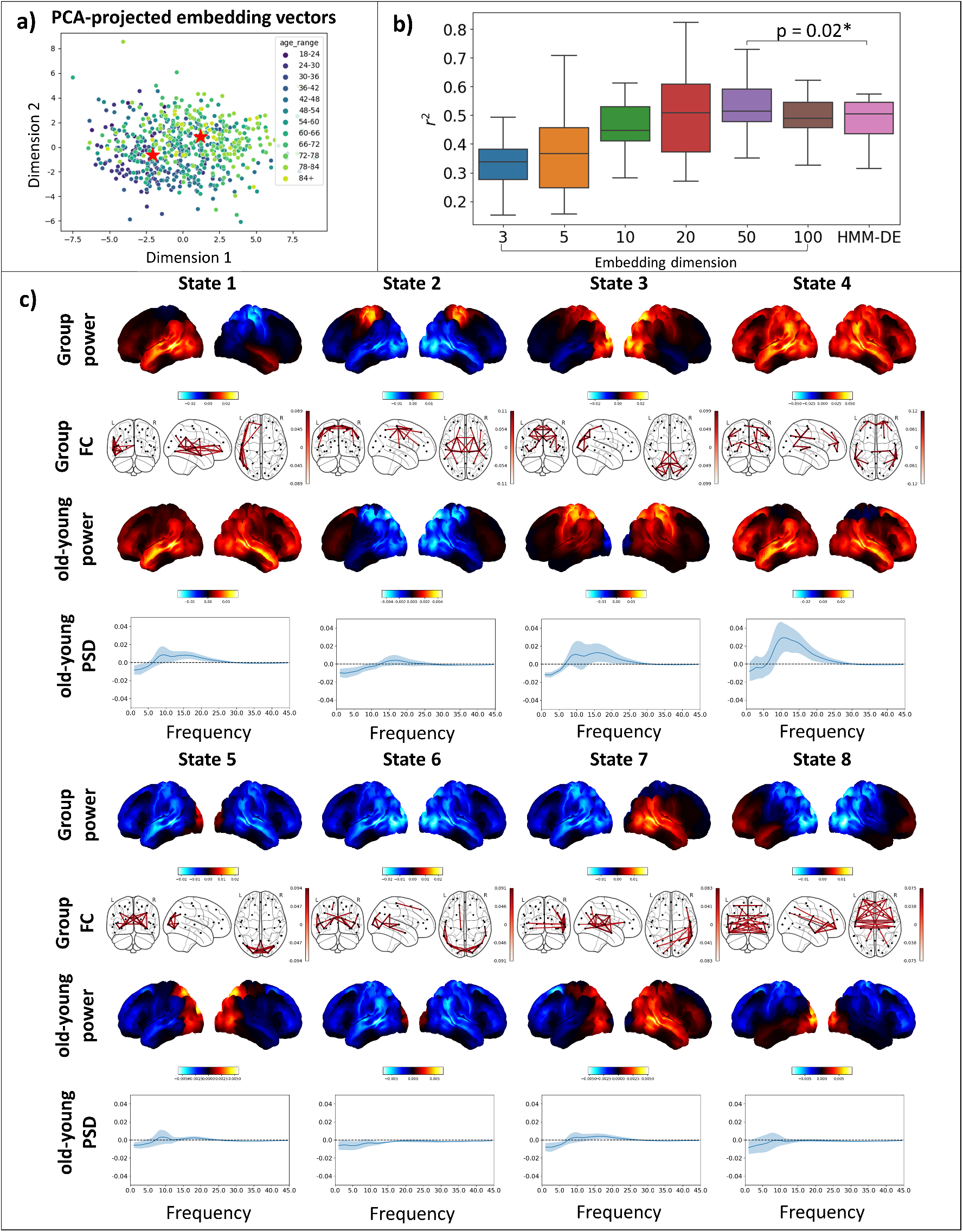
HIVE infers age structure in the population more accurately than dual estimation. *Results obtained using the Cam-CAN dataset*. a) Inferred embedding vectors projected to 2 dimensions with PCA. Darker-coloured points are younger participants and lighter-coloured ones are older participants. The two red stars show centroids of subjects with age < 36 and > 72. b) 20-fold cross validated age prediction accuracy for HIVE with different embedding dimensions and HMM-DE. c) 8 states are inferred by HIVE. Group level power (red areas show above average and blue areas show below average power across states) and FC maps (top 3% edges are plotted) are shown as well as the differences in power (red areas show higher and blue areas show lower power in older subjects than younger subjects) and PSDs (solid lines show the means and shaded areas show one standard deviation across the channels) between old and young subjects.

Centroids of subjects of age groups < 36 and > 72 are used as representatives of young and old subjects. Similar to the analysis in Section 3.2.2, we can find the nearest neighbours (here we choose 20 subjects) for both representatives and get the power maps, PSDs for young and old subjects. We show the results in Figure 10c. We can conclude that HIVE can discover meaningful age patterns in the data.

## 4 Discussion

In this paper we propose the use of embedding vectors as a means of characterising functional networks in different sessions/subjects, similar to how word embedding vectors characterise sematic differences between words in a word dictionary. These embedding vectors are incorporated into a generative model of FC that uses covariance matrices to describe network activity. This can be potentially used in many different variants of brain network models, including static and dynamic FC approaches. Here we apply it to the HMM model and MEG data. In particular, we propose a novel variability encoding block to translate the information encoded in the embedding space to session-specific networks (covariances), which are then fed to the HMM generative model.

Using simulations, we show that during inference the variability encoding block learns multivariate session-specific deviations (Section 3.1). This is an important feature because multivarite session-specific deviations are expected in real data and this model is capable of learning these. Additionally, we also see that the model can accurately infer the pairwise relationships between sessions, i.e. discover subpopulation structure through the embedding vectors. This allows the model to perform more accurate inference than dual estimation on individual sessions with the HMM.

Similar to the approach taken in PROFUMO ([28, 27]), session deviations from the group-level estimates are generated through a Bayesian prior. The additional feature of embedding vectors allows the model to find subpopulations in the group. By training HIVE on the Wakeman-Henson dataset, we observe that inter-subject variability is much greater than inter-session variability (Section 3.2.1). This was also found in fMRI literature ([26]). Although the HIVE inferred covariances have slightly underestimated deviations from the group due to the regularising effect of the Bayesian prior, it is robust to noise in the dataset (Figure 8). Furthermore, the covariances from this approach form better separated clusters due to this de-noising effect compared to the traditional HMM-DE.

In Section 3.2.2, we can observed systematic differences between scanners by training HIVE on data from different sites. The model is trained in an unsupervised fashion and is unaware of the scanners/sites. By finding the centroids of embedding vectors of both datasets and pooling over nearest neighbours, we are able to find the power maps, FC maps and PSD profiles of representatives from both datasets. This allows us to study the systematic differences between scanners. Moreover, HIVE inferred covariances form better-separated clusters than HMM-DE inferred ones based on dataset.

Finally, in Section 3.2.3, by training on a large-scale dataset, we show that the inferred embedding vectors contains age information and there is a clear age gradient. By training classifiers to predict subject age with inferred subject-specific covariances, we see that accuracy increases with embedding dimension and at some point, HIVE inferred features are able to outperform HMM-DE inferred ones. By picking representatives of young and old subjects, and again pooling over information of nearest neighbours, we can acquire brain networks and PSD profiles of young and old brains. The ability to explicitly capture age-related changes could be potentially useful in aging studies.

### Future work

In this paper, we applied embedding vectors to the HMM to model individual sessions. The variability encoding block of session/subject-specific covariance matrices (describing network activity) can be readily adapted to other model-based methods. The simplest example is to set the number of HMM states in HIVE to one, which would make the approach equivalent to a static FC approach while maintaining the power of the embedding vectors to model variability. Another example is DyNeMo ([25]), an alternative method to the HMM, but which makes more flexible assumptions about the dynamics.

HIVE provides a way of learning the underlying population structure, which can be used to regularise the estimates of session-specific measures (e.g. PSD). This could be potentially useful in “boutique” studies that have relatively small numbers of subjects and contain subjects with specific demographics. By training our proposed model on a large-scale dataset and using the learned population structure, individual-level estimates could be better inferred on these small datasets.

Additionally, HIVE can be trained on fMRI data where the number of subjects is much larger, but the number of samples per subject is much smaller, as illustrated by our simulation study (Figure 7). Furthermore, we have shown that HIVE gives better clustering results than dual estimation. A next step will be to investigate the possibility of utilising this advantage in demographics and disease prediction. Lastly, the model provides a natural way to provide subject/session information (e.g. task, age, sex, etc) via the embedding vectors in a hierarchical manner. This could be useful when there is an intrinsic subpopulation structure in the group (e.g. multiple sessions for different subjects and both task and resting-state data are available for the subjects).

### Hyper-parameters

Due to the additional complexity of the model, HIVE has more hyper-parameters than HMM, including the number of layers, number of neurons per layer in the decoders of the variability encoding block. However, in practice, we find that results are very robust to the choice of these hyper-parameters.

The number of states is an important hyper-parameter in HMM that could affect the interpretation of the data. Previous studies show the variational free energy keeps decreasing with increasing number of state ([3]). We hypothesise that this could be due to the inter-session variability in the data (See A.7). Modelling the variability in the data could be a potential solution to this problem.

## 5 Conclusion

We proposed the use of embedding vectors to model individual functional neuroimaging sessions and applied this approach to extend the HMM, giving us HIVE. The variability encoding block explicitly models variability within a population in a principled way. We provide a way to perform efficient inference on the model parameters and the algorithm is readily scalable to large amount of data. With a Bayesian prior, the model pools information across individuals for how they may deviate from the group mean. The embedding vectors allow the model to group together similar data and help the interpretation of sources of variation in a population. This is an important step towards making use of the numerous large-scale datasets collected with different protocols. We believe the above results demonstrate that the proposed model provides a novel perspective in population modelling and in the inference of functional networks.

## 6 Credit authorship contribution statement

RH: Conceptualisation, Methodology, Software, Validation, Formal analysis, Investigation, Data curation, Writing - original draft, Writing - review and editing, Visualisation.

CG: Conceptualisation, Methodology, Software, Data curation, Writing - review and editing. MW: Conceptualisation, Methodology, Data curation, Writing - review and editing, Supervision.

## 7 Acknowledgements

This research was supported by the National Institute for Health Research (NIHR) Oxford Health Biomedical Research Centre. The Wellcome Centre for Integrative Neuroimaging is supported by core funding from the Wellcome Trust (203139/Z/16/Z). RH is supported by the EPSRC Centre for Doctoral Training in Health Data Science (EP/S02428X/1). CG is supported by the Wellcome Trust (215573/Z/19/Z). MW is supported by the Wellcome Trust (106183/Z/14/Z, 215573/Z/19/Z), the New Therapeutics in Alzheimer’s Diseases (NTAD) study supported by UK MRC, the Dementia Platform UK (RG94383/RG89702) and the NIHR Oxford Health Biomedical Research Centre (NIHR203316). The views expressed are those of the author(s) and not necessarily those of the NIHR or the Department of Health and Social Care.

## A Appendix

### A.1 Dual estimation of HMM

Let *γ*_*jt*_ be the state probability of state *j* at time *t*, then the session-specific means 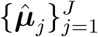 and covariances 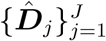 are estimated by

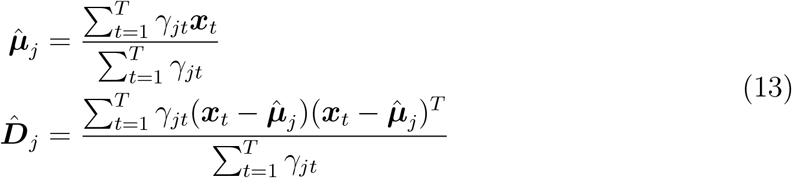

### A.2 Initialisation of variational parameters

We found that the a good initialisation of the variational parameters of the Gamma distributions 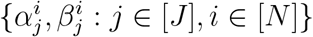 is crucial for the training of HIVE, without which the model can either get stuck in a local minimum or diverge. Here we initialise the shape and rate parameters of the Gamma distributions as follows:

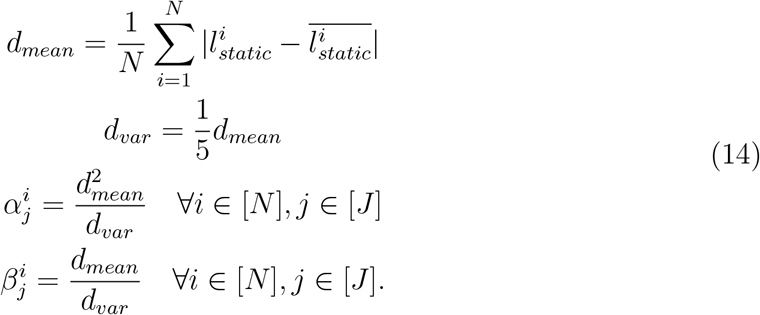

The idea is to estimate the mean deviation magnitude *d*_*mean*_ with the deviation of the Choleksy vectors 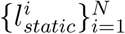 of the static covariance matrices from different sessions and get the parameters so that the resulting Gamma distribution has mean *d*_*mean*_ and variance 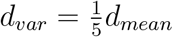, which is what we found to be a good initialisation in practice.

### A.3 Simulating session-specific covariances

Group level covariances ***D***_*j*_ are generated randomly. Each embedding vector is assigned a group uniformly randomly from a set of groups and each embedding vector is generated with a multivariate Gaussian distribution with the same variance but different means depending on the group assignment. Next the encoder and two decoders in Figure 2 are randomly generated linear transformations and the deviation magnitudes 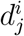 are directly generated by applying the linear transformation on the concatenated embeddings 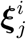. Following the flow chart of Figure 2, we have generated session-specific covariances from different groups of sessions.

### A.4 Outlier in the dataset

We see from Figure 11 that run 3 of subject 8 has a very different static PSD profile from the other runs. In particular the spike at around 26 Hz. Furthermore it has a very different session-pairwise correlation of the static covariances from the other runs.

**Figure 11:**
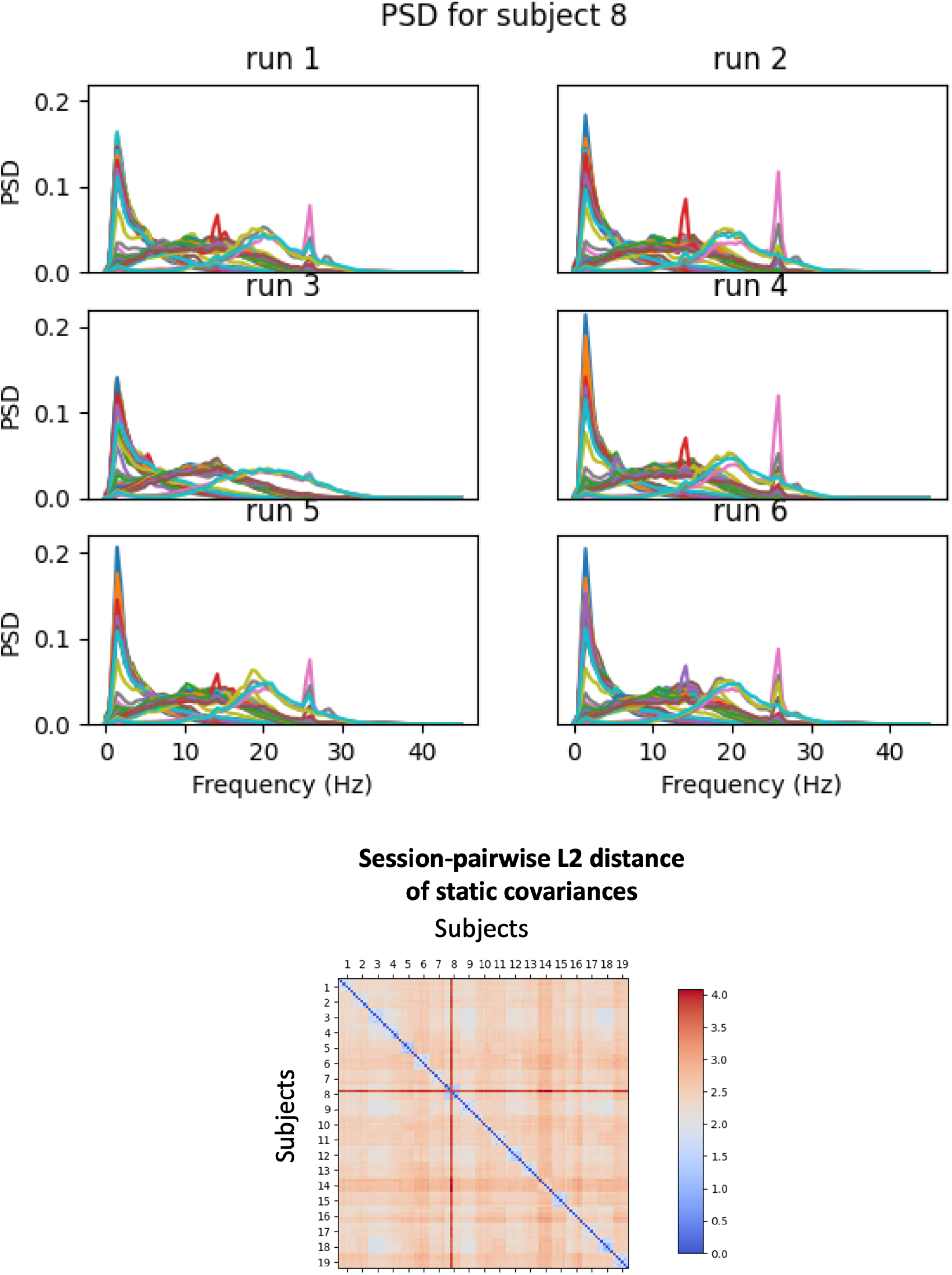
Outlier in the dataset. Top: PSD for the prepared data used for training is plotted for different sessions/runs of subject 8. Bottom: The session-pairwise L2 distance of the static covariances is plotted for the prepared data used for training. The outlier run is highlighted in red.

**Figure 12:**
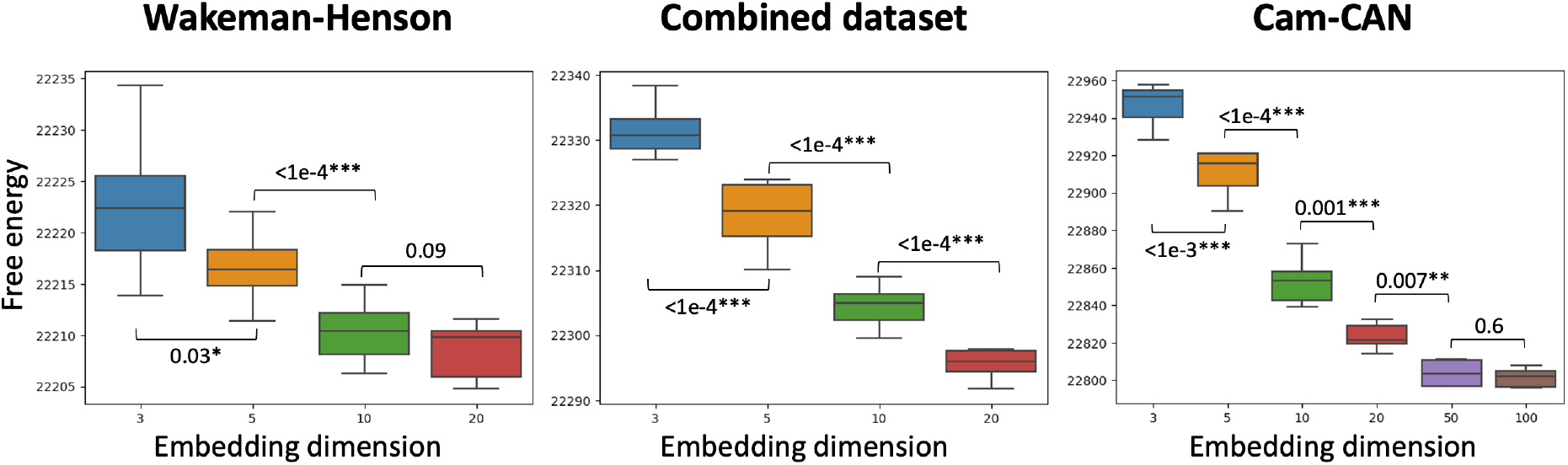
Free energy is used to choose *n*_*ω*_. For each of the real dataset results, different models with different candidate embedding dimensions are trained 10 times independently. The free energy on training data is plotted against embedding dimension. The p-values of the progressive t-tests are plotted between the boxplots.

### A.5 Classifier for predicting age with subject-specific covariances

HMM-DE and HIVE with embedding dimensions 3, 5, 10, 20, 50, 100 are trained on the Cam-CAN dataset. Each of the 7 models is trained 10 times independently and the best model is chosen based on free energy. Next the inferred subject-specific covariances are projected by PCA, whose first few PCs are used as regressors in a ridge regression to predict subject age. The subject age data are separated into 20 folds. For each fold, a classifer is trained on 19 folds and tested on the remaining 1 fold. During training, a 5-fold cross validated grid search is used to select the best dimension of PCA projection and the regularisation strength of the ridge regression. PCA projection dimension is chosen from {5, 20, 50, 100} and regularisation strength is chosen from {10^*i*^ : *i* ∈ {−5, −4, −3, −2, −1, 0, 1, 2, 3}}.

### A.6 Choosing the embedding dimension on the real datasets

We choose *n*_*ω*_ with the procedure described in Section 2.5. For the model trained on the Wakeman-Henson dataset in Section 3.2.1, the decrease in free energy is insignificant when comparing *n*_*ω*_ = 10 and *n*_*ω*_ = 20. Hence for this dataset, we choose *n*_*ω*_ = 10. For the model trained on the combined dataset in Section 3.2.2, the tests are all significant and we choose *n*_*ω*_ = 20. For the model trained on the Cam-CAN dataset in Section 3.2.3, the tests are all significant until *n*_*ω*_ = 50. Hence we choose *n*_*ω*_ = 50.

### A.7 Inter-session variability in data causes ever-decreasing free energy with number of states

We simulate two datasets, one without variability, one with variability between sessions. The ground truth number of states for both datasets is 4. HMM is trained 5 times independently on both datasets and the free energy on training data is plotted in Figure 13. We see that when there is no variability, the free energy decreases with increasing number of states until it plateaus at the ground truth number of states. However, when there is variability, the free energy keeps decreasing with increasing number of states. This is because the model can keep finding more states to explain the inter session variability in the data.

**Figure 13:**
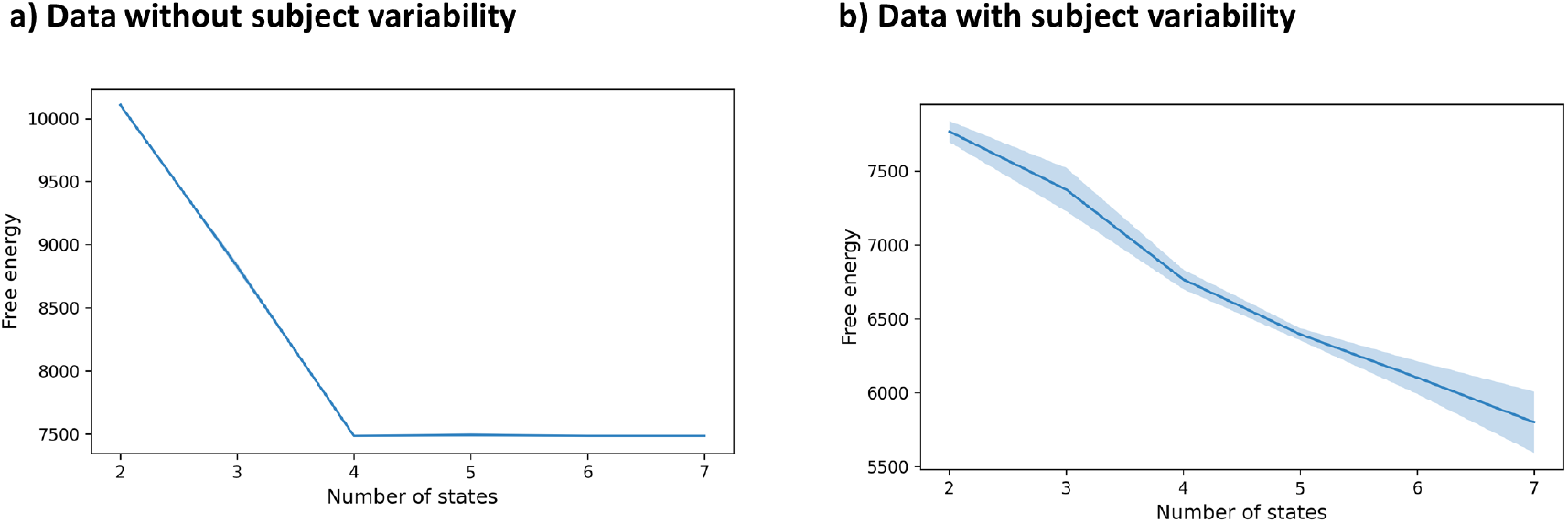
Inter-session variability in the data can cause ever-decreasing free energy in HMM. a) There is no inter-session variability in the data and variational free energy is plotted against number of states. b) There is inter-session variability in the data and variational free energy is plotted against number of states.

